# Blood-brain barrier dysfunction promotes astrocyte senescence through albumin-induced TGFβ signaling activation

**DOI:** 10.1101/2022.04.25.489438

**Authors:** Marcela K. Preininger, Dasha Zaytseva, Jessica May Lin, Daniela Kaufer

## Abstract

Blood-brain barrier dysfunction (BBBD) and accumulation of senescent astrocytes occur during brain aging and contribute to neuroinflammation and disease. Here, we explored the relationship between these two age-related events, hypothesizing that chronic hippocampal exposure to the blood-borne protein serum albumin could induce stress-induced premature senescence (SIPS) in astrocytes via transforming growth factor beta 1 (TGFβ) signaling. We found that one week of albumin exposure significantly increased TGFβ1 and senescence marker expression in cultured rat hippocampal astrocytes. These changes were preventable by pharmacological inhibition of the type I TGFβ receptor (TGFβR) ALK5. To study these effects *in vivo*, we utilized an animal model of BBBD in which albumin was continuously infused into the lateral ventricles of adult mice. Consistent with our *in vitro* results, one week of albumin infusion significantly increased TGFβ signaling activation and the burden of senescent astrocytes in hippocampal tissue. Pharmacological inhibition of TGFβR ALK5 or conditional genetic knockdown of astrocytic TGFβR prior to albumin infusion was sufficient to prevent albumin-induced astrocyte senescence. Together, these results establish a link between TGFβ signaling activation and astrocyte senescence and suggest that prolonged exposure to serum albumin due to BBBD can trigger these phenotypic changes.

## 1. Introduction

Advanced age is a primary risk factor for neurodegenerative disorders that impair cognitive function such as dementia and Alzheimer’s disease (AD) and related dementia (ADRD). Yet, the mechanisms by which advanced age facilitates neuropathology remain elusive. One hypothesis suggests that an age-related increase in physiological and environmental stress induces stress-induced premature senescence (SIPS) in astrocytes, which in turn contributes to the neuronal dysfunction and neuroinflammation associated with age-onset neurodegenerative disorders (Bhat et al., 2012; Bitto et al., 2010; Chinta et al., 2015; Han et al., 2020; Kritsilis et al., 2018; Tan et al., 2014). Cellular senescence is a process that occurs in response to exhausted replication or sublethal levels of cellular stress (McHugh and Gil, 2018). The senescent phenotype is generally characterized by cell cycle arrest, increased lysosomal mass, loss of nuclear integrity, decreased functional capacity, and expression of a senescent-associated secretory phenotype (SASP) (Dodig et al., 2019; Wang and Dreesen, 2018). The prevalence of senescent astrocytes in the brain increases with age, with a significantly higher burden observed in the cortices of patients with AD compared to age-matched controls (Bhat et al., 2012). Recent evidence from mouse models of tau pathology (Bussian et al., 2018) and Parkinson’s Disease (Chinta et al., 2018) support the senescence hypothesis, linking increased astrocyte senescence with neuroinflammation and neurodegeneration. These studies demonstrate that periodically clearing senescent glial cells from the nervous system with targeted interventions can prevent the initiation and progression of disease. While the role of these cells in disease continues to be explored, it is important to identify key physiological and environmental triggers of astrocyte senescence.

One potential physiological trigger of astrocyte senescence is age-associated BBBD and subsequent TGFβ signaling activation. BBBD enables blood-borne proteins such as serum albumin (Bar-Klein et al., 2014; Cacheaux et al., 2009; Senatorov et al., 2019; Weissberg et al., 2015) and fibrinogen (Schachtrup et al., 2010) to extravasate into the brain parenchyma, where they elicit a robust inflammatory response by activating canonical TGFβ signaling. In astrocytes, albumin binds to the type II TGFβR, which dimerizes and subsequently activates the type I TGFβR ALK5 before undergoing receptor-mediated endocytosis. This leads to the phosphorylation of the downstream effector protein SMAD2 and subsequent activation of pro-inflammatory and epileptogenic transcriptional programs (Bar-Klein et al., 2014; Cacheaux et al., 2009; Ivens et al., 2007; Kim et al., 2017). Additionally, albumin-induced TGFβ signaling activation upregulates astrocytic expression of TGFβ and its receptors, resulting in a positive feedback loop that can lead to chronic neurological symptoms (Bar-Klein et al., 2014). Increased BBBD and concomitant albumin leakage has been documented in aging and negatively impacts brain health and function (Farrall and Wardlaw, 2009; Bogdan O. Popescu et al., 2009; Senatorov et al., 2019; Zlokovic et al., 2020). BBBD in the hippocampus, the brain’s center for learning and memory, is especially consequential. BBBD has been observed in the hippocampus of individuals with mild cognitive impairment (MCI) (Montagne et al., 2015; Wang et al., 2006) and occurs before hippocampal atrophy in AD (van de Haar et al., 2016; Van De Haar et al., 2016), suggesting hippocampal BBBD may be critical early step towards neurodegeneration (Montagne et al., 2017).

Notably, studies have demonstrated that TGFβ activation mediates senescence in various cell types *in vitro*, including fibroblasts (Debacq-Chainiaux et al., 2005; Frippiat et al., 2001; Rapisarda et al., 2017), keratinocytes (Vijayachandra et al., 2009), mesenchymal stem cells (MSCs) (Wu et al., 2014), glioblastoma cells (Kumar et al., 2017), cardiomyocytes (Lyu et al., 2018), and lung epithelial cells (Yoon et al., 2005). Also, TGFβ1 as a SASP component was found to be an important mediator of paracrine senescence *in vivo* (Acosta et al., 2013). Thus, we hypothesized that BBBD-induced TGFβ hyperactivation in the brain may play a role in promoting astrocyte senescence. While previous studies have identified the effects of BBBD on astrocyte activation and neurological impairment (Bar-Klein et al., 2014; Cacheaux et al., 2009; Kim et al., 2017; Senatorov et al., 2019), this is the first study to explore its effects on astrocyte senescence. In this study, we sought evidence for a paradigm that integrates BBBD, TGFβ hyperactivation, and senescence—three factors that are consistently associated with age-related neurological disorders. Specifically, we tested the hypothesis that prolonged exposure to the blood serum-derived protein albumin promotes hippocampal astrocyte senescence in a TGFβ-dependent manner.

## 2. Methods

### 2.1. Animal care

All rodent procedures were approved by and performed in accordance with UC Berkeley’s institutional animal care and use committees. Animals were housed under pathogen-free conditions with a 12-hour light and 12-hour dark cycle with food and water available *ad libitum*.

### 2.2 Astrocyte isolation and cell culture

Primary rat hippocampal astrocytes were isolated from P8 Sprague Dawley rat pups (Charles River Laboratories) and cultured under serum-free conditions following an established immunopanning protocol (Foo, 2013). In brief, brain tissues were mechanically and enzymatically dissociated to generate a single-cell suspension, which was incubated successively on antibody-coated plates to subtract microglia, endothelial cells, and oligodendrocyte-lineage cells. After negative selection, astrocytes were positively selected by incubation on an Itgb5-coated plate. Isolated astrocytes were subsequently cultured in a chemically-defined, serum-free medium (50% neurobasal, 50% DMEM, 100 U ml^−1^ penicillin, 100 μg ml^−1^ streptomycin, 1 mM sodium pyruvate, 292 μg ml^−1^ L-glutamine, SATO supplement, 5 μg ml^−1^ N-acetyl cysteine) supplemented with 5 ng ml^−1^ HBEGF as previously described in detail (Foo, 2013).

### 2.3. Immunocytochemistry of cultured astrocytes

Adherent cells were rinsed with cold PBS and fixed with 2% paraformaldehyde solution for 10-15 minutes at room temperature (RT), permeabilized with ice-cold 100% ethanol for 5 min, rinsed again with PBS, and blocked overnight at 4°C with 5% normal goat serum. Cells were incubated for 2 hours at RT with primary antibodies, then rinsed 3 times with PBS to remove excess antibody. Cells were incubated with fluorescently-conjugated secondary antibodies for 1 hour at RT in the dark. Cells were washed 3 more times with PBS. Nuclear counterstaining was performed by adding NucBlue Fixed Cell ReadyProbes Reagent (Thermo Fisher) to the secondary antibody solution. For more information on the antibodies used, refer to **Supplementary Table 1** and **Supplementary Table 2**. Images were acquired and exported using an Echo Revolve (Discover Echo).

### 2.4. Flow cytometry assays

Single cell suspensions of cells harvested from tissue culture were stained using a standard procedure. Briefly, cells were washed with PBS, fixed with 4% paraformaldehyde for 10 minutes, permeabilized with ice-cold 100% EtOH for 3 minutes, and blocked with 3% normal goat serum in PBS for 2 hours at RT. After blocking, cells were incubated with primary antibody or isotype control for 1 hour at RT, washed with PBS, then incubated with fluorescently conjugated secondary antibody for 1 hour at RT in the dark. After incubation with secondary antibody, cells were washed 3 times with PBS. Fluorescent measurements were acquired using a LSR Fortessa flow cytometer (Becton Dickinson) and analyzed with FlowJo Analysis Software (Tree Star).

### 2.5. Gene expression analysis

Total RNA was extracted from astrocytes using Direct-zol RNA MiniPrep kit (Zymo Research) and gene expression was quantified via quantitative reverse transcription PCR (RT-qPCR). First-strand cDNA synthesis was performed from 1 μg of RNA using iScript Reverse Transcription Supermix for RT-qPCR (Bio-Rad). Primers were designed using the NCBI Primer-BLAST tool and supplied by Integrated DNA Technologies. UCSC Genome Browser In-Silico PCR tool was used to confirm the specificity of primer sequences. PCR products were amplified using a CFX96 Real-Time PCR System and threshold cycles were measured using SsoAdvanced Universal SYBR Green Supermix (Bio-Rad). Dissociation curve analysis was performed to verify single products for each reaction. For each sample, reactions were performed in triplicate on 96 well plates (Bio-Rad). Mean threshold cycles were normalized to the *Hprt* housekeeping gene and relative changes in mRNA levels were quantified using the 2^-ΔΔCT^ method (Livak and Schmittgen, 2001). For a list of primer sequences, refer to **Supplementary Table 3**.

### 2.6. Cell culture experiments

Astrocytes were seeded on tissue culture dishes at a density of 10,000 cells per cm^2^ and analyzed for senescence markers after 7 days of treatment. Cells were treated by adding either 10 nM TGFβ1 (PeproTech) or 0.2 mM albumin (Sigma-Aldrich) to fresh culture media every 48 hours. For TGFβ receptor inhibition, cells were pre-treated with either 1 µM SJN2511 (Tocris Biosciences) or DMSO (Sigma-Aldrich) control for 30 minutes prior to albumin treatment.

### 2.7. Senescence-associated β-galactosidase staining

SA-β-Gal staining was performed using a commercially available Senescence β-Galactosidase Staining Kit (Cell Signaling Technologies) according to the manufacturer’s instructions. Quantification was performed by randomly selecting frames and manually counting positive cells using a Zeiss AxioImager microscope (Carl Zeiss) equipped with a Stereo Investigator system (MBF Biosciences). At least 200 cells were counted per sample. Representative images were captured at 20x using an Optronics Microfire A/R color camera.

### 2.8. Osmotic pump implants and IPW-5371 treatment

Osmotic pumps for intracerebroventricular (ICV) delivery of albumin were implanted in adult male C57BL/6 mice under 2% isofluorane anesthesia as previously described (Weissberg et al., 2015, 2011). Using a stereotaxic frame, a hole was drilled through the skull at 0.5 mm caudal, 1 mm lateral to bregma, anterior to the hippocampus. A cannula was placed into the right lateral cerebral ventricle and fixed with surgical glue. Cannulas from Brain Infusion Kit 3 (ALZET) were attached to a Mini-Osmotic Pump Model 2001 (ALZET) filled with 200 µL of either 0.4 mM bovine serum albumin (Sigma-Aldrich) solution or artificial cerebrospinal fluid (aCSF; 129 mM NaCl, 21 mM NaHCO3, 1.25 mM NaH2PO4, 1.8 mM MgSO4, 1.6 mM CaCl2, 3 mM KCl, and 10 mM glucose) and implanted subcutaneously in the right flank. Pumps infused at a rate of 1 µL per hour for 7 days. At day 7, the mice were anesthetized again for pump removal; wounds were closed and fixed with surgical glue. For the IPW-5371-treated cohort, mice received 20 mg/kg of IPW-5371 via intraperitoneal (i.p.) injection daily for 7 days.

### 2.9. Genetic knockdown of astrocytic TGFβ signaling in transgenic mice

Generation and characterization of the triple transgenic TGFβR2 knockdown mice (*a*TGFβR2/KD) used in this study have been previously described in detail (Senatorov et al., 2019). Briefly, the mice were bred to express tamoxifen activated Cre recombinase (CreERT) under the astrocytic promoter glial high affinity glutamate transporter (GLAST) with a floxed exon 4 of tgfbr2 (fl), and a transgenic LacZ reporter gene inhibited by a floxed neomycin cassette. Tamoxifen thus induces activation of astrocytic CreERT resulting in a null TGFβR2 allele and LacZ reporter expression. The inducible Cre/lox system was activated by 5 days of i.p. tamoxifen injection (Sigma Aldrich; 160 mg/kg dissolved in corn oil). GLAST-CreERT; tgfbr2 (fl/+) heterozygotes were used as controls and received the same dosage of tamoxifen. Mice were weighed daily to ensure accurate dosage.

### 2.10. Western blot analysis

Mouse hippocampal tissues were manually homogenized, and protein lysates were extracted using RIPA buffer (50 mM Tris-HCl, 150 mM NaCl, 1% NP-40, 0.5% Sodium deoxycholate, 0.1% SDS) including a protease (Calbiochem) and phosphatase inhibitor cocktail (Roche). Protein samples were run under reducing conditions. 20 µg of protein lysate was mixed with 4x Laemmli buffer (Bio-Rad) containing 5% 2-mercaptoethanol (Sigma) and fractionated by SDS-PAGE using the Mini-PROTEAN Tetra System (Bio-Rad) and Mini-PROTEAN TGX Precast gels (4-20%, 15-well comb, 15µL/well). Each gel was loaded with 7 µL of Chameleon Duo pre-stained protein ladder (LI-COR) for reference. Following separation, samples were transferred to a 0.45 µm polyvinylidene fluoride (PVDF) membrane (Bio-Rad). Membranes were blocked for 1 hr at RT with Intercept TBS blocking buffer (LI-COR) and incubated overnight at 4°C with primary antibodies. Membranes were then washed 3 times for 10 mins with TBST (0.05% Tween-20 in TBS) and incubated with secondary antibodies in blocking buffer for 1 hr at RT in the dark. Membranes were washed 3 times for 10 mins with TBST and visualized using the Odyssey DLx Imaging System (LI-COR). Fluorescence analysis was performed in Image J (NIH). Blot values were normalized to GAPDH internal control and mean relative changes in protein levels were normalized to the control group. For details about the antibodies used, refer to **Supplementary Table 1** and **Supplementary Table 2**.

### 2.11. Mouse tissue sampling and immunohistochemistry

Mice were anesthetized with Euthasol euthanasia solution and transcardially perfused with ice cold, heparinized (10 units/mL) physiological saline for 10 minutes followed by 4% paraformaldehyde (PFA) in PBS. Mouse brains were removed and post-fixed in 4% PFA in PBS for 24 hours at 4°C, then cryoprotected by saturating with 30% sucrose in PBS. Brains were embedded in Tissue Plus O.C.T. compound (Scigen) and frozen at -80°C. Brains were later sliced on a CryoStar NX70 cryostat into 20 µm coronal sections and mounted onto charged slides for immunostaining. Prior to staining, slides were treated for antigen retrieval by incubating them in 10 mM citrate buffer (pH 6.0) with 0.05% Tween-20 at 70°C for 20 minutes. For added permeabilization, slides were incubated with 0.01% Triton X-100 in Tris-buffered saline (TBS) for 10 minutes at RT. To prevent nonspecific binding, slides were incubated with blocking buffer composed of 10% normal donkey serum in TBS with 0.05% Tween-20 for 1 hour at RT. After initial blocking, tissues were incubated overnight with primary antibodies diluted 1:200 in blocking buffer. The next day, slides were washed 3 times in TBS and subsequently incubated with fluorescent conjugated secondary antibodies diluted 1:800 in blocking buffer in the dark for 2 hours at RT. Nuclear counterstaining was performed by adding NucBlue Fixed Cell ReadyProbes Reagent (Invitrogen) to the secondary antibody solution. Slides were washed 3 times in TBS and mounted with ProLong Diamond (Invitrogen). For details about the antibodies used, refer to **Supplementary Table 1** and **Supplementary Table 2**.

### 2.12. Lamin B1 quantification

Immunostained sections were imaged with a Nikon TiE inverted confocal microscope using a 100x oil objective. Z-stacks were acquired using 1 µm steps for a total of 20 steps per region using NIS Elements imaging software (Nikon). Fluorescence intensity was quantified using ImageJ (NIH) as previously described (Chinta et al., 2018). Nuclei of GFAP+ and GFAP-cells were segmented within each Z-stack and mean gray value was measured in the channel corresponding to Lamin B1. At least 5 GFAP+ astrocytes and 5 GFAP-cells were sampled per section and at least 2 hippocampal sections were sampled per animal.

### 2.13. Statistical analysis

Statistics and graphs were performed using GraphPad Prism 8 software. For the *in vitro* time course experiment, a two-way ANOVA was used followed by Tukey’s multiple comparisons test. For all other experiments, a one-way ANOVA was used followed by Dunnett’s post-hoc test to compare treatment groups to the control group when a main effect was detected. For correlations, two-tailed linear regressions were performed to determine Person’s r and p-value. Significance thresholds were set at *p < 0.05, **p < 0.01, ***p < 0.001.

## 3. Results

### 3.1. Albumin exposure activates TGFβ signaling in cultured hippocampal astrocytes

To test the effects of albumin exposure on astrocytes *in vitro*, hippocampal astrocytes were isolated from postnatal day 8 rat pups, a time at which albumin is ostensibly excluded from the developing rat cortex (Bento-Abreu et al., 2008). The hippocampus was selected for analysis because it is especially vulnerable to age-associated BBBD (Pelegri et al., 2007). To avoid the confounding effects of serum proteins, astrocytes were isolated and maintained in chemically-defined, serum-free media using a previously established protocol (Foo et al., 2011). After isolation, cultures were expanded to passage 3 before being used for experiments. Cultures were 97% positive for the astrocytic marker glial fibrillary acidic protein (GFAP) as assessed via flow cytometry, and displayed typical astrocytic morphology with multiple branch-like processes when imaged via immunocytochemistry (**Fig. 1A**)

**Figure 1.**
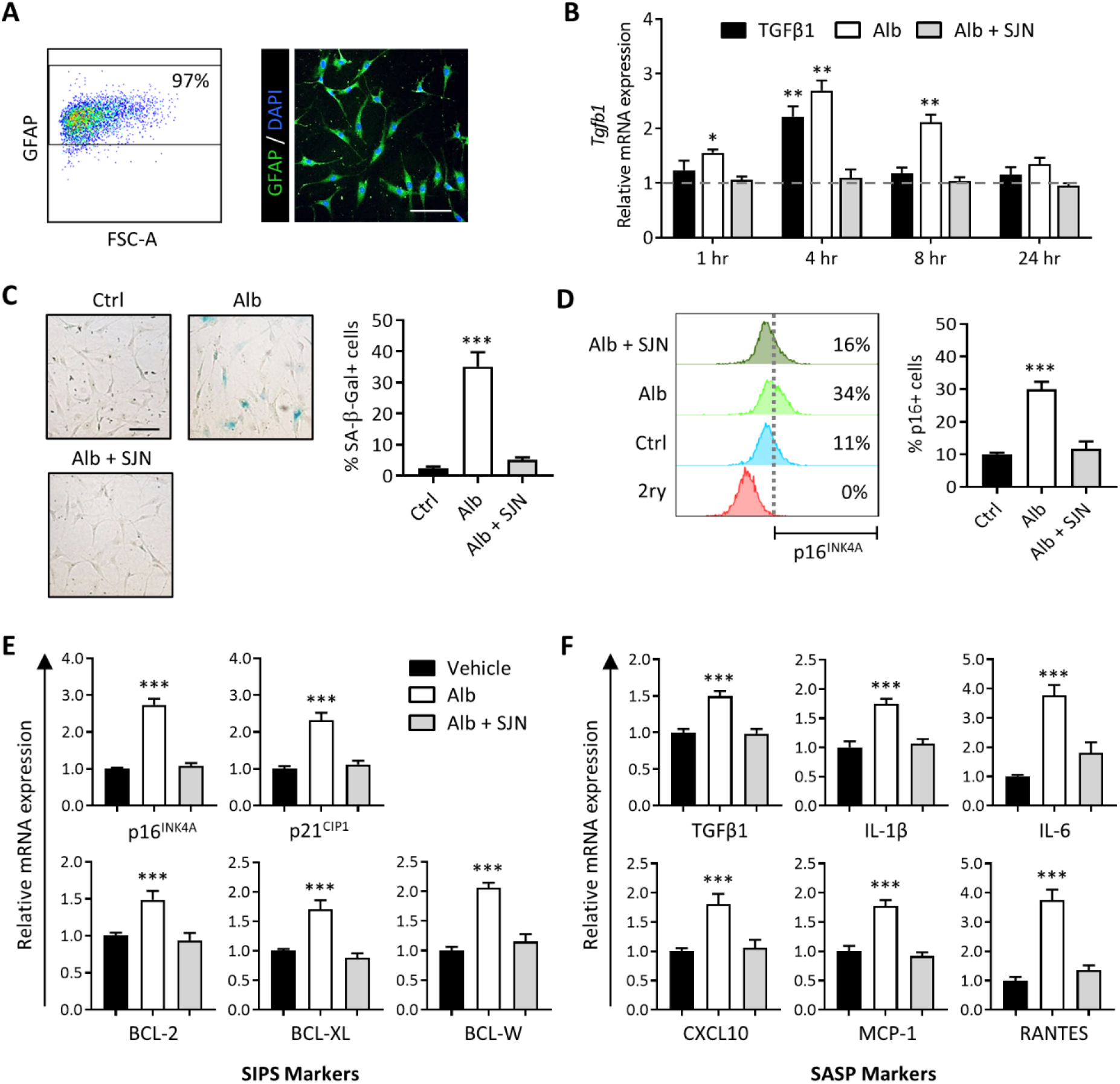
Albumin-induced TGFβ signaling activation induces astrocyte senescence in cultured hippocampal astrocytes. (A) Representative dot plot and microscopy image show astrocytes isolated from P8 rat hippocampi were 97% GFAP positive and exhibit typical morphology with multiple processes. (B) Bar graph shows relative *Tgfb1* mRNA expression levels against baseline (horizontal dotted line). Astrocytes exhibited a significant increase in *Tgfb1* transcription upon exposure to albumin, reaching maximum *Tgfb1* mRNA levels at 4 hours after exposure. Albumin-induced *Tgfb1* transcription was blocked by TGFβR inhibition with SJN2511. (C) Left: representative brightfield images show 7 days of albumin exposure induces senescence-associated morphological changes in astrocytes and increases SA-β-Gal expression. Right: Quantification via cell counting reveals a significant increase in the percentage of SA-β-Gal positive cells in the albumin-treated cells compared to control, which was blocked by TGFβR inhibition with SJN2511. (D) Left: histograms depict the results of one representative flow cytometry experiment to illustrate the gating strategy applied for determining p16^INK4A^ positive cells. The bottom (red) histogram represents the isotype control used to set the fluorescence detection threshold (vertical dotted line). Cells with fluorescent values above the threshold were considered positive for p16^INK4A^. Right: Summary data of cell counts from all experiments show that treatment with albumin induced significant increases in the percentage of p16^INK4A^ positive cells compared to control. SJN2511 treatment prevented the albumin-induced increase in p16^INK4A^ positive cells. (E) Gene expression analysis after 7 days of albumin exposure revealed increases in mRNA levels of genes encoding markers of stress-induced premature senescence (SIPS) and (F) genes encoding components of the senescence-associated secretory phenotype (SASP). Albumin-induced upregulation of these genes was prevented by TGFβR inhibition with SJN2511. Scale bar: 100 µm. Bar graphs depict mean ± SEM of 3 independent experiments. Asterisks denote a significant difference from baseline *p <0.05, **p<0.01, ***p<0.001.

Previous studies have demonstrated that albumin binds to the type II TGFβR and activates canonical type I TGFβR ALK5, leading to the phosphorylation of the downstream effector protein SMAD2. Consequently, albumin exposure leads to increased levels of *Tgfb1* mRNA in astrocytes (Bar-Klein et al., 2014; Cacheaux et al., 2009; Ivens et al., 2007; Ralay Ranaivo et al., 2010). To determine the kinetics and dose-dependence of this response *in vitro*, astrocytes were treated with 0.2 mM of albumin and harvested after 1, 4, 8, and 24 hours of exposure for gene expression analysis via RT-qPCR. The albumin concentration used in this study is consistent with previous *in vitro* studies investigating its signaling effects on primary astrocytes and derived from *in vivo* transient hippocampal albumin measurements following BBBD (Cacheaux et al., 2009; Frigerio et al., 2012; Ralay Ranaivo et al., 2012). Treatment with 10 nM of TGFβ1 under the same conditions was also included as a previously validated positive control for TGFβ signaling activation in astrocytes (Bar-Klein et al., 2014). Astrocytes exhibited increased *Tgfb1* transcription in response to albumin treatment, reaching peak *Tgfb1* mRNA levels at 4 hours after exposure to albumin or TGFβ1 and remained significantly elevated in the albumin condition for at least 8 hours. Importantly, treatment with SJN2511, a selective small molecule TGFβR ALK5 inhibitor, blocked albumin induced upregulation of *Tgfb1* across all timepoints, with no significant difference observed from baseline (**Fig. 1B**).

### 3.2 Albumin-induced TGFβ signaling increases expression of senescence markers in cultured astrocytes

To test whether albumin induces astrocyte senescence, we exposed cultures to 0.2 mM albumin for 7 days and analyzed them for cellular markers of senescence. A 7 day timepoint was selected in accordance with other *in vitro* protocols where prolonged exposure to a non-lethal stressor induced senescence in primary cells (Chinta et al., 2018; Kumar et al., 2017; Yu et al., 2017). One common feature of senescent cells is increased lysosomal mass, which is measured by the detection of senescence associated β-galactosidase (SA-β-Gal) (Dimri et al., 1995). At day 7, albumin-treated astrocytes exhibited SA-β-Gal expression and flat morphology consistent with the senescent phenotype, whereas the control group exhibited no morphological changes (**Fig. 1C**). In the control group, only ∼3% of the cells were SA-β-Gal positive at day 7. In albumin-treated cultures, a significant increase in the percentage of SA-β-Gal positive cells compared to control was observed, with a mean of 35.1% positive cells (**Fig. 1C**). TGFβR inhibition with SJN2511 prevented the albumin-induced increase in astrocyte senescence as detected by SA-β-Gal expression, limiting the percent of SA-β-Gal positive cells to a mean of 4.2% at day 7 with no significant difference from control (**Fig. 1C)**.

Along with lysosomal changes, senescent cells exhibit changes in pathways related to cell cycle control and apoptosis that contribute to their survival after stressful conditions. Another marker of cellular senescence is the expression of p16^INK4A^, a cyclin dependent kinase inhibitor that is upregulated upon cell cycle exit (Serrano et al., 1997). To test for expression of p16^INK4A^, cells were assessed by flow cytometry at day 7. A detection gate was set to include single cells only, and the fluorescence threshold for p16^INK4A^ positivity was defined using a secondary-only negative control. In untreated control cells, a mean of ∼10% of cells were detected as p16^INK4A^ positive. Consistent with the SA-β-Gal assay results, 7 days of albumin exposure significantly increased the percentage of p16^INK4A^ positive cells compared to control, with a mean of approximately 30% p16^INK4A^ positive cells detected in the albumin-treated condition (**Fig. 1D**). TGFβR inhibition with SJN2511 prevented the albumin-induced increase in astrocyte senescence. An average of 11.7% cells were detected as p16^INK4A^ positive in the SJN2511 treatment group, which was not significantly different from the control group (**Fig. 1D**).

### 3.3. Albumin-induced TGFβ signaling increases expression of senescence-associated genes in cultured astrocytes

In addition to p16^INK4A^, senescent cells often exhibit activation of the p21^CIP1^ pathway, associated with growth arrest (Herbig et al., 2004), and BCL family proteins, which provide resistance to apoptosis (Yosef et al., 2016). Gene expression analysis via RT-qPCR after 7 days albumin exposure revealed increased mRNA levels of cyclin dependent kinase inhibitors 2a (*Cdkn2a*; p16^INK4A^) and 1a (*Cdkn1a*; p21^CIP1^) and the apoptosis regulators *Bcl2* (BCL-2), *Bcl2l1* (BCL-XL), and *Bcl2l2* (BCL-W) compared to control (**Fig. 1E**). Albumin-induced upregulation of these genes was prevented by pharmacological TGFβR inhibition with SJN2511 (**Fig. 1E**). These data support the hypothesis that prolonged albumin exposure promotes expression of senescence-associated genes, including cell cycle and apoptosis inhibitors, in cultured hippocampal astrocytes in a TGFβ-dependent manner.

Another feature of senescent cells is SASP expression, so we performed gene expression analysis on a subset of SASP factors that are associated with AD pathogenesis. Gene expression analysis via RT-qPCR after 7 days albumin exposure revealed increases in mRNA levels of genes encoding transforming growth factor beta 1 (*Tgfb1*; TGFβ1), interleukin 1 beta (*Il1b*; IL-1β), interleukin 6 (*Il6*; IL-6), C-X-C motif chemokine ligand 10 (*Cxcl10*; CXCL10), monocyte chemoattractant protein-1 (*Ccl2*; MCP-1), and C-C motif chemokine ligand 5 (*Ccl5;* RANTES) (**Fig. 1F)**. The albumin-induced activation of all measured SASP genes was prevented by TGFβR inhibition, with no significant differences observed between controls and SJN2511 treated cells (**Fig. 1F)**. These data show that prolonged albumin exposure in cultured hippocampal astrocytes induces changes in gene expression of key components of both the SIPS and proinflammatory SASP phenotypes, and that these changes are dependent on activation of the TGFβR ALK5.

### 3.4. *In vivo* brain albumin infusion decreases nuclear membrane integrity in astrocytes, which is preventable by pharmacological TGFβR inhibition

To test whether exposure of the brain environment to albumin *in vivo* also leads to an accumulation of senescent astrocytes, we used an established mouse model of BBBD to simulate blood protein extravasation in the brain (Milikovsky et al., 2019; Senatorov et al., 2019; Weissberg et al., 2015). In this model, small osmotic pumps were implanted to continuously infuse 0.4 mM albumin or artificial cerebrospinal fluid (aCSF) directly into the right lateral ventricles of adult mice via for 7 days. The 0.4 mM albumin concentration was selected for intracerebroventricular (ICV) administration as it corresponds to normal albumin concentration in blood serum (Merlot et al., 2014) and robustly triggers TGFβ signaling in rodents (Milikovsky et al., 2019; Senatorov et al., 2019; Weissberg et al., 2015). aCSF is a biologically inert fluid in the brain parenchyma and is used as a sham control in this model.

To further evaluate the role of TGFβ signaling in albumin-induced senescence, we tested the efficacy of IPW-5371, a small molecule TGFβR ALK5 inhibitor, in wild type adult mice receiving albumin infusions. IPW-5371 is an experimental drug with a promising clinical profile, including the ability to cross the BBB and a favorable PK/PD profile that enables once-per-day dosing (Rabender et al., 2016). IPW-5371 has been previously validated as a TGFβR ALK5 inhibitor in the context of BBBD in aged brains, where it was shown to significantly reduce levels of phosphorylated SMAD2 in the hippocampi of IPW-5371-treated mice compared to control mice (Senatorov et al., 2019).

In the IPW-5371 treatment cohort, mice received daily intraperitoneal injections (20 mg/kg) of IPW-5371 for 7 days following pump implant. On day 7, the pumps were removed, and the mice were allowed to recover. One week after pump removal, the mice were sacrificed, and the brains were harvested and prepared for immunohistochemistry (IHC) **(Fig. 2A**). Since commercially available p16^INK4A^ antibodies are not suitable for IHC in mice, we used nuclear Lamin B1 quantification to assess astrocyte senescence in mouse hippocampal tissue. Reduced Lamin B1 expression is detectable via IHC and is an established marker of cellular senescence (Chinta et al., 2018; Freund et al., 2012). Only the right hippocampus ipsilateral to the pump infusion was assayed for each section, specifically the hippocampal area of the dentate gyrus delimited by the granule cell layer and the end of the CA3 cell layer. In mice that received 7 days of albumin infusion, hippocampal astrocytes (GFAP+ cells) had significantly reduced Lamin B1 expression compared to control mice infused with aCSF (**Fig. 2B, C**). In hippocampal non-astrocytes (GFAP-), no significant difference was detected in albumin-infused mouse tissues compared to control tissues (**Fig. 2B, D**). Importantly, albumin-induced astrocyte senescence was prevented by TGFβR inhibition with IPW-5371, with no significant differences observed between controls and IPW-5371-treated mice **(Fig. 2B, C**). These findings suggest that albumin-induced astrocyte senescence is mediated by TGFβ signaling activation, and that astrocytes preferentially undergo senescence via this mechanism.

**Figure 2.**
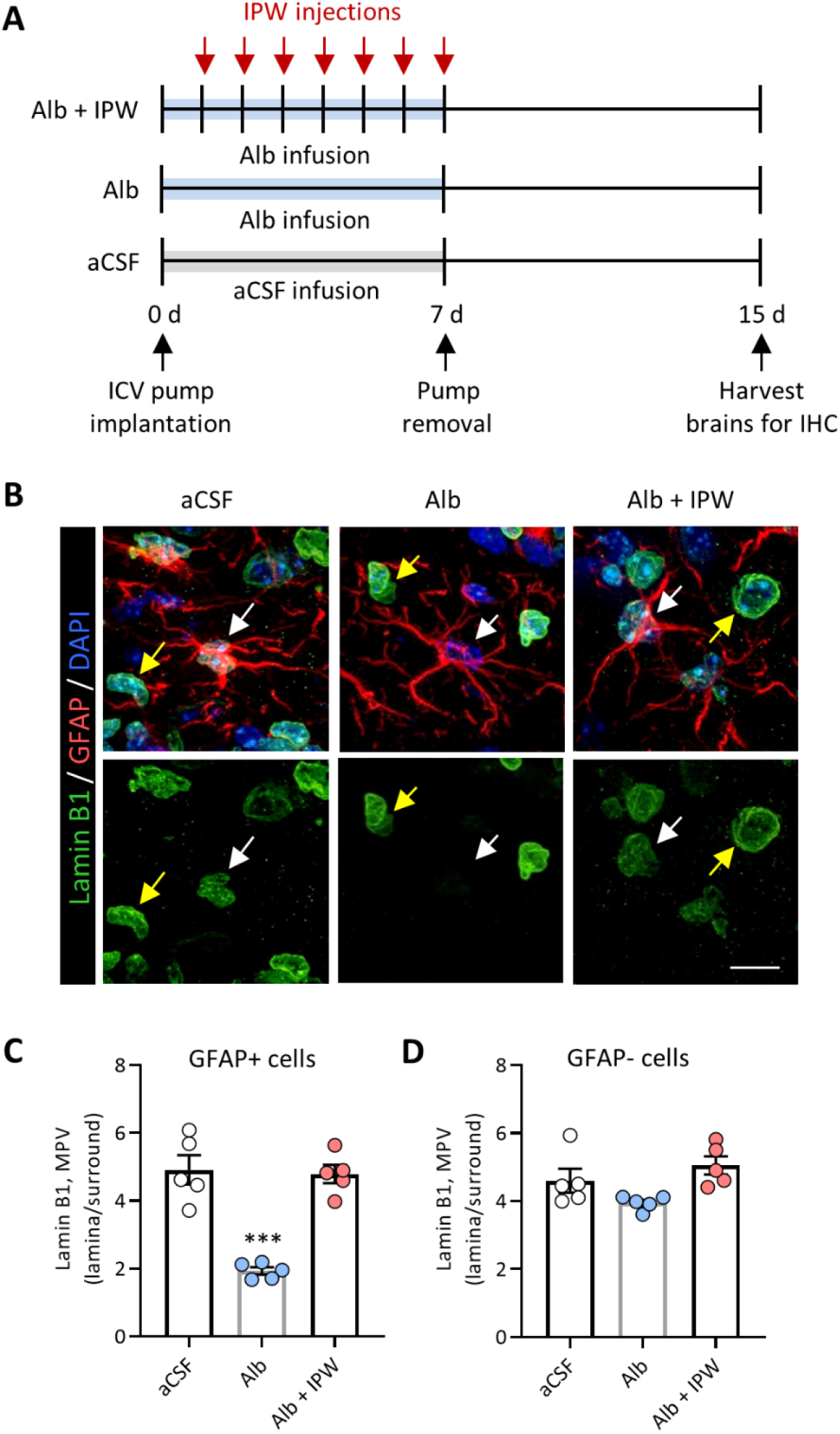
Pharmacological TGFβR inhibition prevents albumin-induced astrocyte senescence in a mouse model of BBBD. (A) Schematic of experimental design whereby mice were infused for 7 days with either aCSF or albumin before pump removal and 7 days of recovery. The control group received aCSF infusion (n=5), another group received albumin infusion only (n=5), and the treatment group received albumin infusion plus 7 days of daily IPW-5371 injections (n=5). Whole brains were extracted 7 days after pump removal and prepared for IHC. (B) Representative confocal Z-stack images of hippocampal sections immunostained for Lamin B1 (green), GFAP (red), and DAPI (blue). White arrows denote GFAP+ cells and yellow arrows denote GFAP-cells. Images acquired at 100x magnification. (C) Quantification of Lamin B1 expression using mean pixel values (MPV) show that in albumin treated mice, GFAP+ astrocytes, but not GFAP-cells, have significantly decreased Lamin B1 levels compared to controls. Pharmacological TGFβR inhibition with IPW-5371 prevents albumin induced Lamin B1 deficits in GFAP+ astrocytes. Scale bar: 10 µm. Bar graphs depict mean ± SEM of each experimental group. Asterisks denote a significant difference from control *p <0.05, **p<0.01, ***p<0.001.

### 3.5. ICV albumin infusion increases hippocampal expression of senescence markers, which is preventable by astrocytic TGFβR knockdown

To test the causal role of TGFβ signaling in albumin-induced astrocyte senescence, we used a previously established transgenic mouse line (*a*TGFβR2/KD) expressing inducible Cre recombinase under the astrocytic-specific glial high-affinity glutamate transporter (GLAST) promoter, which enables conditional knockdown (KD) of the floxed (fl) type II TGFβ receptor (TGFβR2) specifically in astrocytes upon treatment with tamoxifen (Senatorov et al., 2019). Tamoxifen induced efficient recombination in ∼40% of hippocampal astrocytes and TGFβR protein levels were significantly ablated in hippocampal tissue samples, indicating successful genetic knockdown (Senatorov et al., 2019). Selective knockdown of TGFβR2 alone is sufficient to inhibit TGFβ signaling since membrane dimerization of type II and type I TGFβRs is required for phosphorylation of the TGFβR ALK5 intracellular kinase domain and subsequent downstream signaling. Mice that were heterozygous for floxed TGFβR2 (fl/+) were used as controls. Using the same model as before, albumin or aCSF was infused directly into the right lateral ventricles of adult mice via osmotic pumps for 7 days. After 7 days of infusion, the pumps were removed, and the mice were allowed to recover. Two months after pump removal, the mice were sacrificed, and right hippocampi were harvested for protein extraction (**Fig. 3A**).

**Figure 3.**
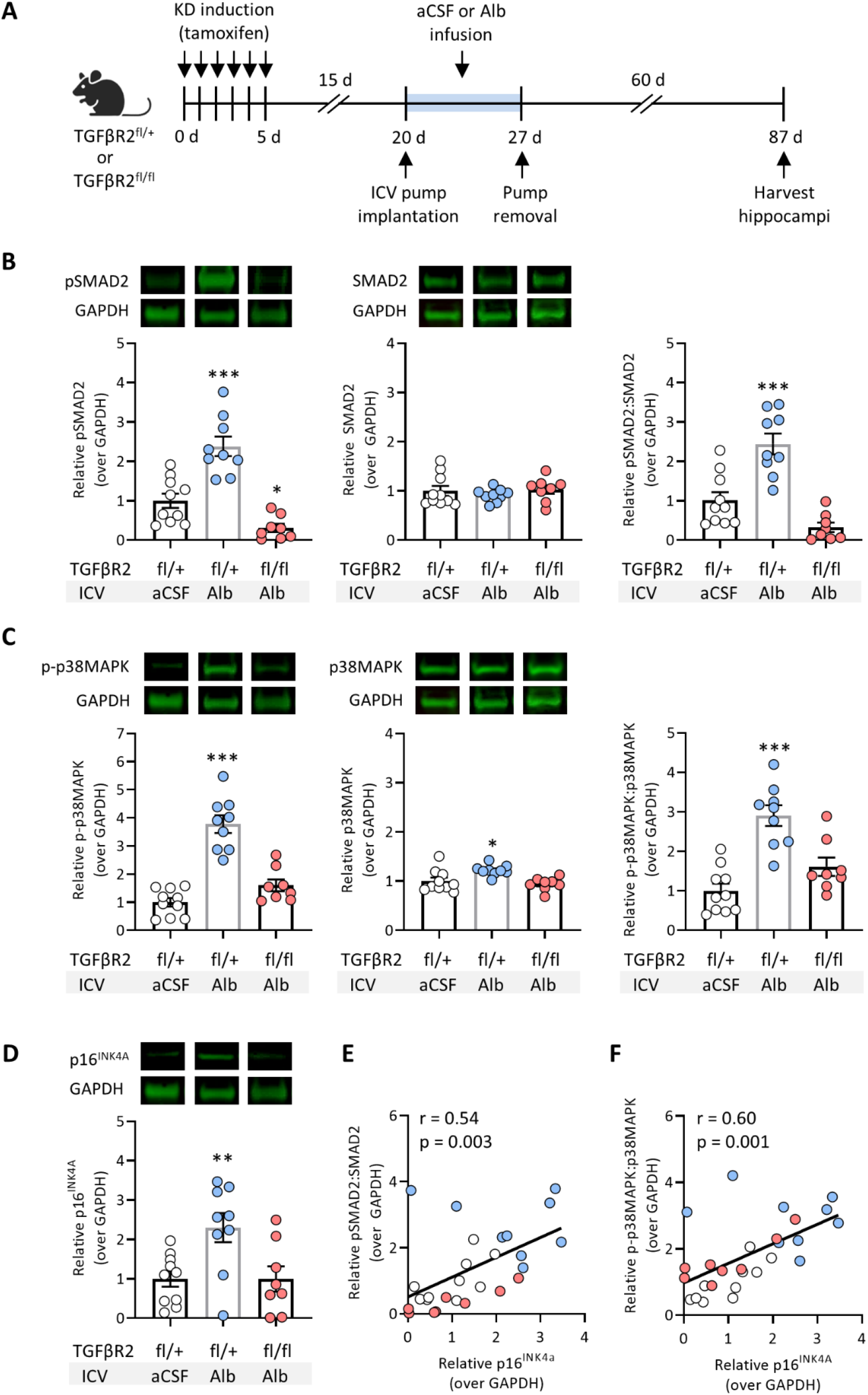
Astrocytic TGFβR knockdown prevents albumin-induced hippocampal senescence in a mouse model of BBBD. (A) Schematic of experimental design whereby ICV osmotic pumps were implanted into the right ventricles of adult transgenic mice. Mice heterozygous for floxed TGFβR2 (fl/+) were infused with aCSF (n=10) or albumin (n = 9), and mice with homozygous induced KD of TGFβR2 (fl/fl) were infused with albumin (n=8) for 7 days before pump removal and recovery. Hippocampal tissue was harvested 60 days after pump removal. Representative Western blot images and bar graphs show protein expression levels normalized to GAPDH of (B) pSMAD2, SMAD2, and the ratio of pSMAD2:SMAD2, (C) p-p38MAPK, p38MAPK, and the ratio of p-p38MAPK:p38MAPK, and (D) p16^INK4A^ in aCSF and albumin-treated mice. Regression analysis revealed positive correlations between the protein expression levels of (E) p16^INK4A^ and the ratio of pSMAD2:SMAD2, and (F) p16^INK4A^ and the ratio of p-p38MAPK:p38MAPK. Bar graphs depict mean ± SD of each experimental group. Asterisks denote a significant difference from control *p <0.05, **p<0.01, ***p<0.001.

TGFβR2 fl/+ control mice that received ICV albumin infusion had significantly higher levels of TGFβ signaling compared to mice that received aCSF infusion, as measured by levels of phosphorylated SMAD2 (pSMAD2) and the ratio of pSMAD2 to unphosphorylated SMAD2 (**Fig. 3B**). To explore an alternative senescence marker, we assessed activation of the p38 mitogen-activated protein kinase (p38MAPK) pathway, as indicated by its phosphorylated state (p-p38MAPK). Activation of p38MAPK is believed to play a causal role in cellular senescence following persistent low-level stresses (Iwasa et al., 2003) and is essential for SASP expression (Freund et al., 2011). Therefore, p-p38MAPK has been used as an additional indicator of senescence in astrocytes (Bhat et al., 2012; Mombach et al., 2015) and neurons (Jurk et al., 2012). In albumin-treated TGFβR2 fl/+ mice, the levels of p38MAPK, p-p38MAPK, and the ratio of p-p38MAPK to p38MAPK were significantly higher compared to aCSF-treated controls (**Fig. 3C**). Notably, albumin infusion also significantly increased p16^INK4A^ expression in TGFβR2 fl/+ mice compared to controls (**Fig. 3D**). Mice with homozygous induced KD of TGFβR2 (fl/fl) in astrocytes were protected against albumin-induced increases in TGFβ signaling and senescence marker expression (**Fig. 3B-D**). These results show that genetic KD of astrocytic TGFβ signaling is sufficient to prevent albumin-induced senescence in these cells.

Regression analysis revealed there were positive correlations between the pSMAD2:SMAD2 ratio and p16^INK4A^ expression (r = 0.54, p = 0.003) (**Fig. 3E**) and the p-p38MAPK:p38MAPK ratio and p16^INK4A^ expression (r = 0.60, p = 0.001) (**Fig. 3F**) across all groups. Together, these data support a relationship between albumin-induced astrocytic TG-β signaling and elevated senescence pathways in the brain.

## 4. Discussion

Senescent astrocytes are present in aging brain tissue and play a role in neurological disease progression, but the specific triggers of age-related senescence remain elusive. In this study, we sought to test the hypothesis that the blood-derived protein albumin, which accumulates in the aging brain due to BBBD, can induce astrocyte senescence through TGFβ hyperactivation. Using a serum-free, *in vitro* model, we discovered that one week of serum albumin exposure induced expression of senescence and SASP markers in isolated hippocampal astrocytes, and that pharmacological inhibition of the TGFβR ALK5 was able to prevent these phenotypic changes. Using an *in vivo* model of BBBD, we found that one week of continuous albumin infusion into the lateral ventricles of mice preferentially caused senescence in astrocytes, and that pharmacological TGFβR inhibition with IPW-5371 was able to prevent albumin-induced astrocyte senescence in mice. Using the same model, we found that 7 days of albumin infusion in control mice caused increased TGFβ signaling (pSMAD2) and senescence marker expression (p16^INK4A^ and p-p38MAPK) in mouse hippocampi two months after pump removal. Senescence marker expression was TGFβ-dependent since selective knockdown of astrocytic TGFβR prior to albumin infusion was sufficient to prevent these increases. Together, these results establish a link between TGFβ hyperactivation and astrocyte senescence and suggest that prolonged albumin exposure due to BBBD can underlie these phenotypic changes.

The albumin-induced increase in transcripts encoding the AD-associated SASP factors IL-1β, IL-6, CXCL10, MCP-1, and RANTES supports the link between senescent astrocytes and neurodegeneration. For instance, IL-1β and IL-6 are both potent stimuli for leukocyte recruitment to the brain and have been implicated in neuroinflammation and neurofibrillary pathology in AD (Bradburn et al., 2018b; Campbell, 1998; Campbell et al., 1993; Li et al., 2003; Sheng et al., 2001). Chemokines serve a similar function in recruiting immune cells to the CNS during AD (Altstiel and Sperber, 1991). For instance, CXCL10 signals through the CXCR3 receptor to promote monocyte activation and migration (Liu et al., 2011). CXCL10 is found in high concentrations in the brains of AD patients and activation of the CXCL10/CXCR3 axis has been shown to mediate development of AD-like pathology in mouse models (Krauthausen et al., 2015). In humans, peripheral levels of CXCL10 are elevated in older adults and negatively associated with working memory performance (Bradburn et al., 2018a). Overexpression of MCP-1 has been shown to enhance microgliosis, facilitate amyloid plaque formation, and accelerate cognitive decline in rodent models of AD (Kiyota et al., 2009; Yamamoto et al., 2005). Although its role in neuropathology is still being investigated, RANTES is upregulated in the cerebral microcirculation of AD patients and is also suspected to influence neuroinflammation (Tripathy et al., 2010).

Previous studies have demonstrated that senescent astrocytes have substantially reduced functional capacity compared to their normal counterparts. For example, senescent astrocytes exhibited impaired would healing ability, phagocytic uptake, metabolic function, and neuroprotective capacity (Bang et al., 2021; Pertusa et al., 2007). Among their numerous functions in the adult brain, astrocytes are required for BBB maintenance (Heithoff et al., 2021). Therefore, it’s possible that initial BBBD-induced astrocyte senescence establishes a positive feedback loop that results in progressive neurological decline. In this scenario, the accumulation of functionally impaired senescent astrocytes and secretion of pro-inflammatory SASP factors results in more BBBD and thus more TGFβ-induced astrocyte senescence. Further studies should investigate how this process is regulated and explore potential strategies to prevent its onset.

While this study has identified blood-derived serum albumin as a potential physiological trigger of TGFβ signaling activation and astrocyte senescence, it is unlikely to be the only activator of these pathways in the aging brain. Previous reports have identified additional age-related triggers of astrocyte senescence such as oxidative stress (Bitto et al., 2010), proteotoxic stress (Bitto et al., 2010), and amyloid beta (Aβ) oligomers (Bhat et al., 2012). Interestingly, TGFβ signaling is also implicated in these age-related senescence mechanisms, making our results consistent with past findings. In the case of oxidative stress, TGFβ increases the production of reactive oxygen species (ROS) by impairing mitochondrial function and inducing NADPH oxidases; TGFβ also suppresses the synthesis of antioxidant enzymes such as glutathione in various cell types, causing redox imbalance and promoting further oxidative stress (Krstić et al., 2015; Liu and Desai, 2015; Liu and Gaston Pravia, 2010). In the case of Aβ pathology, TGFβ overexpression increases astrocytic Aβ generation in transgenic mice (Lesné et al., 2003; Wyss-Coray et al., 1997, 1995), and TGFβ levels are elevated in cortical astrocytes surrounding Aβ plaques in AD patients (Apelt and Schliebs, 2001; van der Wal et al., 1993).

Given the salience of TGFβ signaling activation in response to physiological triggers of senescence, this pathway should be explored as an attractive therapeutic target in the prevention of age-related neuropathology. Future studies should seek to elucidate the context in which various triggers promote senescence during aging and the extent of their respective contributions.

## Abbreviations

BBB: blood-brain barrier
BBBD: blood-brain barrier dysfunction
TGFβ: transforming growth factor beta 1
ALK: activin like receptor
AD: Alzheimer’s disease
ICV: intracerebroventricular

## Acknowledgements

We thank the Cancer Research Laboratory Flow Cytometry Facility and the Molecular Imaging Center at UC Berkeley for the use of their instruments. We also thank Dr. Vladimir Senatorov, Dr. Oscar Vazquez, Vaidehi Ghandi, and Bridget Ma for their experimental assistance, and Dr. Barry Hart and Innovation Pathways Inc. for supplying IPW-5371. This research was supported by a Bakar Foundation Fellowship (D.K.), the Archer Foundation Award (D.K.), Borstein Family Foundation award (D.K), NSF GRFP fellowship (M.K.P.), and NIH T32 fellowship (GM 098218; M.K.P.).

## Author contributions

Marcela Preininger: Conceptualization, Investigation, Formal analysis, Writing -Original draft; Dasha Zaytseva: Investigation, Validation, Formal analysis; Jessica May Lin: Investigation, Validation, Formal analysis; Daniela Kaufer: Supervision, Resources, Funding Acquisition, Writing – Review & Editing

