## Supplemental Material for "Blood-brain barrier dysfunction promotes astrocyte senescence through albumin-induced TGFβ signaling activation"

to

**Supplementary Table 1.** Primary Antibodies

| Target | Isotype | Supplier | Catalog # | Dilution | Application |
|---|---|---|---|---|---|
| GFAP | Goat IgG | Abcam | ab53554 | 1:500 | FACS, ICC, IHC |
| Lamin B1 | Rabbit IgG | Abcam | ab16048 | 1:200 | IHC |
| $p16^{INK4a}$ | Mouse IgG | Abcam | ab54210 | 1:500 | FACS |
| $p16^{INK4a}$ | Rabbit IgG | Assay Biotech | C0285 | 1:500 | WB |
| SMAD2 | Rabbit IgG | Cell Signaling | 5339S | 1:1000 | WB |
| pSMAD2 | Rabbit IgG | Millipore Sigma | AB3849-I | 1:1000 | WB |
| p38 MAPK | Rabbit IgG | Cell Signaling | 9212S | 1:1000 | WB |
| p-p38 MAPK | Rabbit IgG | Cell Signaling | 4511S | 1:1000 | WB |
| GAPDH | Rabbit IgG | Cell Signaling | 2118S | 1:1000 | WB |

FACS = flow cytometry; ICC = immunocytochemistry; IHC = immunohistochemistry; WB = Western blot

**Supplementary Table 2.** Secondary Antibodies

| Conjugate | Host and Isotype | Supplier | Catalog # | Dilution | Application |
|---|---|---|---|---|---|
| Alexa 488 | Donkey anti-rabbit IgG | Thermo Fisher | A-21206 | 1:800 | IHC |
| Alexa 594 | Donkey anti-goat IgG | Thermo Fisher | A-11058 | 1:800 | IHC |
| Alexa 488 | Goat anti-mouse IgG | Thermo Fisher | A-32723 | 1:500 | FACS, ICC |
| IRDye 800CW | Goat anti-rabbit IgG | LI-COR | 926-32211 | 1:10,000 | WB |

FACS = flow cytometry; ICC = immunocytochemistry; IHC = immunohistochemistry; WB = Western blot

**Supplementary Table 3.** Gene Primers for RT-qPCR

| Gene | Primer Sequence |
|---|---|
| *Hprt* | 5’ – TCAGTCAACGGGGGACATAAA – 3’<br>3’ – GGGGCTGTACTGCTTAACCAG – 5’ |
| *Tgfb1* | 5’ – CAACCCAGGTCCTTCCTAAA – 3’<br>3’ – GGAGAGCCCTGGATACCAAC – 5’ |
| *Cdkn2a* | 5’ – AATCTCCGCGAGGAAAGC – 3’<br>3’ – GTCTGCAGCGGACTCCAT – 5’ |
| *Cdkn1a* | 5’ – ATCACCAGGATTGGACATGG – 3’<br>3’ – GGTGTCAGAGTCTAGGGGA – 5’ |
| *Bcl2l1* | 5’ – GCTGCATTGTTCCCGTAGAG – 3’<br>3’ – GTTGGATGGCCACCTATCTG – 5’ |
| *Bcl2* | 5’ – GGTCTTCAGAGACAGCCAGG – 3’<br>3’ – GATCCAGGATAACGGAGGCT – 5’ |

| | |
|---|---|
| *Bcl2l2* | 5' – TCTAGTGGCTGACTTTGTAGGC – 3'<br>3' – GAAACCTGGGTGAAGCGTTG – 5' |
| *Ccl2* | 5' – GCATCTGCCCTAAGGTCTTCA – 3'<br>3' – GTGGAAAAGGTAGTGGATGCATT – 5' |
| *Il1b* | 5' – CACAGCAGCACATCAACAAG – 3'<br>3' – GTGCTCATGTCCTCATCCTG – 5' |
| *Ccl5* | 5' – CCCTCACCATCATCCTCACT – 3'<br>3' – TCCTTCGAGTGACAAACACG – 5' |
| *Ccl20* | 5' – TGTACGAGAGGCAACAGTCG – 3'<br>3' – TCTGCTCTTCCTTGCTTTGG – 5' |
| *Il6* | 5' – GCTACCAAACTGGATATAATCAGGA – 3'<br>3' – CCAGGTAGCTATGGTACTCCAGAA – 5' |
